# Retigabine and gabapentin restore channel function and neuronal firing of an epilepsy-associated dominant-negative *KCNQ5* variant

**DOI:** 10.1101/2023.03.24.534091

**Authors:** Johanna Krüger, Holger Lerche

## Abstract

**Objective:** *KCNQ5* encodes the voltage-gated potassium channel K_V_7.5, a member of the K_V_7 channel family, which conducts the M-current. This current was shown to be a potent regulator of neuronal excitability by mediating the medium and slow afterhyperpolarization. Recently, we have identified five loss-of-function variants in *KCNQ5* in patients with genetic generalized epilepsy. Using the most severe dominant-negative variant p.(Arg359Cys) (R359C), we set out to investigate pharmacological therapeutic intervention by K_V_7 channel openers on channel function and neuronal firing.

**Methods:** Whole-cell patch clamp recordings were conducted in human embryonic kidney cells to investigate the immediate effect of retigabine, gabapentin and intracellular application of zinc on the R359C variant in absence and presence of K_V_7.5-WT subunits. Transfected primary hippocampal cultures were used to examine the effect of R359C on neuronal firing and whether this effect could be reversed by drug application.

**Results:** Retigabine and gabapentin both increased R359C-derived K^+^ current density and M-current amplitudes in both homomeric and heteromeric mutant K_V_7.5 channels. Retigabine was most effective in restoring K^+^ currents. Ten µM retigabine was sufficient to reach the level of WT currents without retigabine, whereas 100 µM of gabapentin showed less than half of this effect and application of 50 µM zinc only significantly increased M-current amplitude in heteromeric channels. Overexpression of K_V_7.5-WT potently inhibited neuronal firing by increasing the M-current, and medium afterhyperpolarization, whereas R359C overexpression had the opposite effect. All three aforementioned drugs reversed the effect of R359C reducing firing to nearly normal levels at high current injections.

**Significance:** Our study shows that a dominant-negative complete loss-of-function variant in K_V_7.5 leads to largely increased neuronal firing indicating a neuronal hyperexcitability. K_V_7 channel openers, such as retigabine or gabapentin, could be treatment options for otherwise pharmacoresistant epilepsy patients carrying loss-of-function variants in *KCNQ5*.

## Introduction

*KCNQ5* encodes the α-subunit of K_V_7.5, a delayed rectifier potassium channel that is widely expressed in brain, skeletal muscle, and blood vessels (Lerche et al. 2000, Schroeder et al. 2000, Brueggemann et al. 2007). These subunits can assemble either as homo- or heterotetramers with K_V_7.3 subunits to form channels that – together with K_V_7.2 – generate the M-current (Lerche et al. 2000, Schroeder et al. 2000). This current governs neuronal excitability by suppressing repeated firing through K^+^ outward current during membrane depolarization (Brown and Adams 1980). However, channel opening is not only controlled by changes in membrane potential but also by activation of muscarinic receptors triggering a G-protein coupled cascade resulting in depletion of phosphatidylinositol-4,5-bisphosphate (PIP_2_) from the membrane causing an inability for the channel to open. This enables precise intrinsic control and manipulation of repetitive neuronal firing by acetylcholine release inhibiting the M-current and thus enabling repetitive firing (Delmas and Brown 2005). Consequently, dominant-negative loss-of-function (dnLOF) results in decreased medium and slow afterhyperpolarization currents in the hippocampal CA3 region (Tzingounis et al. 2010), modified hippocampal network synchronization, and reduced synaptic inhibition (Fidzinski et al. 2015). Furthermore, two dnLOF mouse models have been published exhibiting handling- and temperature-induced absence and myoclonic seizures in adult but not in young animals (Wei et al. 2022). This is in contrast to the early onset phenotypes observed in patients with LOF variants in KCNQ2 and KCNQ3 that cause self-limited familial neonatal epilepsy (SLFNE) or developmental and epileptic encephalopathy (DEE; Singh et al. 1998, Charlier et al. 1998, Weckhuysen et al. 2012). Pathogenic gain-of-function (GOF) variants were described in severe intellectual disability (ID) with or without seizures, and DEE (Lehman et al. 2017, Nappi et al. 2022, Wei et al. 2022). Additionally, two truncating variants in less severely affected individuals and an intragenic duplication in a patient with absence seizures, migraine, and mild ID have been reported, all resulting in a haploinsufficiency (Wei et al. 2022, Rosti et al. 2019). Recently, we identified five heterozygous LOF variants in KCNQ5 in patients with genetic generalized epilepsy (GGE). These patients develop epilepsy at a median age of 7±1.6 years with absence seizures being the predominant seizure type of onset in younger individuals, whereas an onset in adolescence is usually characterized by myoclonic or generalized tonic-clonic seizures. Half of these individuals also present with a mild to moderate ID. Of these patients, only one became seizure-free without medication and three are pharmacoresistant (Krüger et al. 2022). Knowing the underlying cause of the disease enables us to explore personalized treatments. However, the first choice K_V_7 channel opener, retigabine (RTG), which activates K_V_7 channels by probably binding to their activation gate and pore region (Wuttke et al. 2005), was withdrawn from the market. While improved versions of RTG are in clinical trials and currently unavailable to patients, other lines of potential treatments have to be explored. Recently, gabapentin (GBP) was proposed to activate K_V_7 channels with the highest sensitivity in homomeric K_V_7.5 channels (Manville and Abbott 2018) and has been successfully used to treat a KCNQ2 LOF patient (Soldovieri et al. 2020). Furthermore, intracellular zinc application has been shown to activate K_V_7 channels by reducing their dependence on PIP_2_ (Gao et al. 2017) and interfering with their inhibition through calmodulin (Yang et al. 2023). Here, we set out to investigate the effect of RTG, GBP and intracellular zinc application on wildtype (WT) and dnLOF K_V_7.5 channels to investigate their potential as treatment options in both transfected human embryonic kidney (HEK) cells as well as in primary hippocampal neuronal cultures transfected with a complete dnLOF K_V_7.5 variant (R359C). In addition, we were able to show how this variant alters neuronal firing in the absence of any drug and could thus confirm K_V_7.5 as a key contributor to the neuronal M-current.

## Materials and Methods

### Human embryonic kidney (HEK) cell culture and transfection

HEK293 cells were grown in Dulbecco’s modified Eagle medium (DMEM; Gibco) containing 10% (v/v) fetal calf serum (FCS; PAN Biotech) and were cultured at 37°C in a 5 % CO_2_ humidified atmosphere. Cells were plated in 35 mm dishes and transfected 24-48 h later using the same protocol as in Krüger et al. 2022. In brief, a Lipofectamine 3000 (Invitrogen) protocol was used according to the manufacturer using 2 µg of either wildtype (WT) or variant DNA resulting in homomeric channel expression. For co-transfection of both subunits 1µg of WT pcDNA3.1-KCNQ5-P2A-tRFP and 1µg pcDNA3.1-KCNQ5-P2A-eGFP DNA of the variant were applied resulting in heteromeric expression, while 1µg of pcDNA3.1-KCNQ5-P2A-tRFP-WT served as the positive control. To obtain untransfected but treated cells as a negative control, the DNA was replaced by water in both conditions. After 24-48 h, electrophysiological experiments were conducted.

### Primary neuronal culture and transfection

Animal protocols for primary cell culture were approved by the local Animal Care and Use Committee (Regierungspraesidium Tuebingen, Tuebingen, Germany). Pregnant C57BL/6NCrl mice were sacrificed on embryonic day 18 (E18) and the embryos were quickly decapitated and removed as described before (Hedrich et al. 2014). The hippocampi were extracted from the harvested embryonic brains and washed thrice with ice-cold magnesium- and calcium-free Hanks’ balanced salt solution (HBSS; Gibco) before applying 2.5 % trypsin and subsequent incubation at 37°C for 14 min. The hippocampi were then rinsed with DMEM supplemented with FCS (PAN Biotech), and penicillin/streptomycin (PAN Biotech). Neurons were dissociated by trituration, subsequent filtering through a cell strainer and then seeded onto poly-D-lysine (Sigma-Aldrich) coated 13 mm coverslips, and cultured in DMEM supplemented with FCS, L-glutamine (PAN Biotech) and penicillin/streptomycin (PAN Biotech) at 37 °C in a 5 % CO_2_ humidified atmosphere. After 4 h, the medium was replaced by Neurobasal culture medium (Gibco) supplemented with B27 (Gibco), L-glutamine and penicillin/streptomycin.

Cultured hippocampal neurons were transfected using Optifect (Invitrogen) according to the manufacturer’s instructions at day *in vitro* (DIV) 5. Concisely, 4µl Optifect reagent was added to 100 µl Opti-MEM (Gibco) and incubated at room temperature (RT) for 5 min. Ensuing, 1 µl water (control) or 1 µg of either WT or variant DNA was added to the mix and incubated at RT for 20 min before being added dropwise to the well. Electrophysiological recordings were performed 72 h after transfection.

### Electrophysiology

An Axopatch 200B amplifier, a Digidata 1550B digitizer (Axon Instruments), and pCLAMP 10.7 or 11.1 data acquisition software (Molecular Devices) were used for standard whole-cell voltage and current clamp recordings at RT. For voltage-clamp protocols, a pre-pulse protocol (-P/4) was used to automatically subtract capacitive and leakage currents. Series resistance was regularly monitored and compensated at approximately 85 %. For voltage-clamp recordings, currents were filtered at 1 kHz and digitized at 5 kHz, while for current-clamp recordings currents were low-pass filtered at 10 kHz and sampled at 100 kHz. The bath solution contained (in mM): 160 NaCl, 2.5 KCl, 1 MgCl_2_, 2 CaCl_2_, 10 HEPES (pH 7.4 adjusted with NaOH) (Gao et al. 2017). Retigabine (10µM; Sigma-Aldrich) and gabapentin (100µM; Sigma-Aldrich) were applied directly to the bath solution shortly before recording in non-saturating concentrations that have been used and proven to be efficacious previously (Soldovieri et al. 2020). Borosilicate glass pipettes had a final tip resistance of 3-6 MΩ and were filled with pipette solution containing (in mM): 120 K-acetate, 35 KCl, 5 NaCl, 3 Na_2_ATP, 10 HEPES (pH 7.3 adjusted with KOH). For intracellular application of zinc, 50µM ZnCl_2_ were added to the pipette solution. The liquid junction potential was calculated as 10.8 mV and not corrected. An inverted microscope (Axio-Vert.A1, Zeiss) was used to visualize transfected cells.

### Patch clamp protocols and data analysis

For voltage-clamp recordings in HEK cells, a step protocol to determine the current density and activation curves was applied as described in Krüger et al. 2022. To measure the M-current, cells were held at 0 mV and hyperpolarized for 1 s to -60 mV. Recordings in hippocampal neurons were conducted in voltage-clamp mode to record the medium afterhyperpolarization current (I_mAHP_) and M-current; all other recordings in neurons were conducted in current-clamp mode. To determine the M-current, the protocol mentioned above was changed to a holding potential of -30 mV. The AHP current was evoked by depolarizing the membrane to +45 mV for 100 ms from a holding potential of -55 mV. The mAHP was measured as the immediate peak after stepping back to -55 mV relative to baseline (Kim et al. 2016). To obtain the input resistance, a series of currents ranging from -10 to -110 pA was injected in 10 pA increments, plotted against the corresponding steady state voltage response, and the slope of a fitted linear regression was calculated. Action potential (AP) trains were evoked by injection of a series of currents ranging from -50 to +300 pA in 25 pA increments for 800 ms. APs were only considered as such when the voltage peak amplitude surpassed 0 mV. Single APs were evoked by injecting the minimum amount of current for 5 ms to evoke an action potential. Data analysis was performed using Clampfit 10.7 and 11.1 (Molecular Devices) and Microsoft Excel (Microsoft Corporation). Statistical analyses were performed using GraphPad Prism 9.5.1 (GraphPad Software). Data is tested for normal distribution and shown as mean ± standard error of the mean (SEM). Normally distributed data was evaluated using either one-way analysis of variance (ANOVA) with Dunnett’s *post hoc* test or two-way ANOVA with Tukey’s *post hoc* test to compare the effect of the different drugs on the WT and variant channels. If data was not normally distributed a Kruskal-Wallis test with Benjamini, Krieger and Yekutieli *post hoc* test was used. For all statistical tests p<0.05 was considered significant.

## Results

The K_V_7.5 R359C variant has been previously reported by us in a family suffering from GGE with some individuals additionally being diagnosed with mild to moderate ID. Functional studies in Chinese hamster ovarian cells revealed a severe dnLOF effect showing almost no current following co-expression of K_V_7.5- or K_V_7.3-WT subunits (Krüger et al. 2022). As one of these individuals is pharmacoresistant to many anti-seizure medications, we set out to test three known K_V_7 channel openers on their efficacy on the variant to provide K_V_7.5 LOF patients with a potential treatment option. Besides retigabine (RTG) and gabapentin (GBP), intracellular application of ZnCl_2_ was investigated as it was proposed that elevated intracellular zinc levels lock K_V_7 channels in the open state by making them independent of PIP_2_ binding and insensitive to inhibition by calmodulin binding (Gao et al. 2017, Yang et al. 2023). As residue R359 is involved in PIP_2_ binding and its mutation likely abolishes an interaction hence locking the channel in a closed state (Krüger et al. 2022), we also wanted to investigate whether zinc application would be able to overcome the PIP_2_ binding defect and enable the mutant channels to open.

### Pharmacological characterization of the K_V_7.5 R359C variant in HEK cells

To investigate the effect of RTG, GBP and intracellular zinc application on K_V_7.5-WT and mutant channels in absence of other channels, HEK cells were transiently transfected as described above. Homomeric WT channels showed robust K^+^ currents whereas the R359C homomeric channels displayed a complete LOF as described previously (Figure 1A; Krüger et al. 2022). Application of 10 µM RTG significantly increased WT peak current density as well as M-current amplitudes, and both current density and M-current level of homomeric R359C channels was restored to that of WT channels under control conditions (Figure 1A, B, E, K, N). RTG application shifted the activation curve to more hyperpolarized potentials, however it was only significant in the R359C compared to untreated WT cells. Furthermore, RTG significantly flattened the slope of the activation curve for the R359C variant (Figure 1H; Supplemental Table 1). Application of 100 µM GBP did not have a significant effect on WT channels. However, peak current densities and M-current amplitudes were increased in cells expressing R359C subunits by GBP, yet currents were still significantly lower as compared to WT channels (Figure 1A, C, F, L, O). Activation curves were not significantly altered by drug application (Figure 1I). Intracellular zinc application did not significantly increase peak current density or M-current in neither WT nor R359C homomeric channels, (Figure 1A, D, G, M, P). Activation curves did not show significant alteration compared to the WT (Figure 1J).

**Figure 1.**
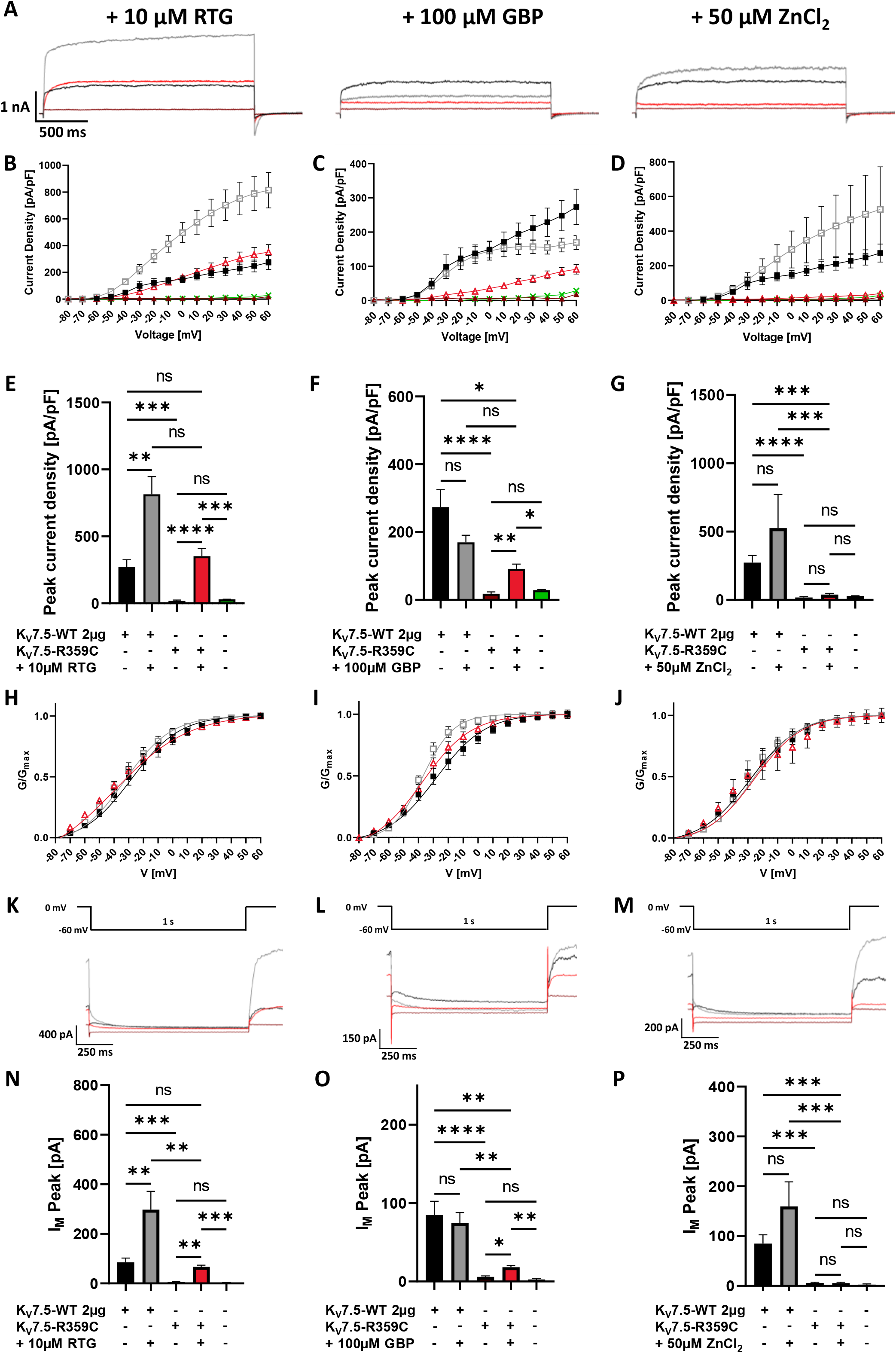
Pharmacological characterization of homomeric K_V_7.5-WT and -R359C channels in HEK cells. HEK cells were transfected with either KCNQ5-WT or KCNQ5-R359C DNA and recorded under either RTG (left panel), GBP (middle panel) or ZnCl_2_ application (right panel). (**A**) Representative K^+^ current traces at +60 mV for K_V_7.5-WT (control black, drug treated grey) and K_V_7.5-R359C (control maroon, drug treated red). (**B-D**) Peak K+ currents were normalized to cell capacitances and plotted versus voltage. R359C (filled marron triangles) currents are significantly decreased as compared to WT (filled black squares) currents showing no difference to CTRL cells (green crosses). RTG significantly increased WT currents (grey outlined squares) and increased R359C peak current density (red outlined triangles) back to WT levels. GBP had no significant effect on WT currents but significantly increased mutant channel currents, though they were still significantly decreased as compared to WT. Intracellular application of ZnCl_2_ did not significantly increase currents in the WT or mutant channels. (**E-G**) Maximum current peak density comparison at +60 mV of WT (black) and R359C (maroon) to drug treatment (grey or red, respectively). (**H-J**) Voltage dependent activation curves. Normalized tail currents were fit with a Boltzmann function. No such relationship could be established for the R359C as currents were too small. RTG significantly left-shifted R359C as compared to WT. (**K-M**) Representative M-current traces and voltage protocol. (**N-P**) M-current peak was significantly increased in WT and R359C by RTG application, GBP only increased R359C peak currents, and ZnCl_2_ did cause significant change. Shown are means ± SEM (**E-G** and **N-P**). * *p* < 0.05, ** *p* < 0.01; *** *p* < 0.001; **** *p* < 0.0001 (one-way ANOVA with Dunnett’s *post hoc* test or Kruskal-Wallis test Benjamini, Krieger and Yekutieli’s *post hoc* test). Supplemental Table 1 provides exact numbers and statistical analysis.

Experiments using co-expression of both WT and R359C channels yielded comparable results as homomeric expression, yet zinc application significantly increased M-current in heteromeric R359C channels (see Supplemental Figure 1 and Table 1).

### Functional characterization of K_V_7.5-WT and -R359C variant in transfected primary cultured hippocampal mouse neurons

In the next set of experiments, we investigated the effect of the dnLOF of R359C versus a WT overexpression on neuronal properties in transfected primary murine hippocampal neuronal cultures. Whole-cell patch-clamp recordings were performed in current clamp mode to determine intrinsic neuronal parameters, action potential characteristics and firing properties, as well as in voltage clamp mode to investigate the M-current. As the R359C variant showed a nearly complete dnLOF in HEK cells, it was assumed it would have the same effect on the endogenous K_V_7.5 and K_V_7.3 subunits, thus creating a condition close to a heterozygous patient. We transfected another batch of neurons with KCNQ5-WT cDNA to induce an overexpression of the channel to mimic a gain of function. Untransfected neurons were used as controls (CTRL). The firing frequency of R359C-transfected neurons was largely increased as compared to untransfected control conditions, whereas WT overexpressing neurons showed a severe decrease in firing to the point that only few neurons would fire a single action potential (13/35) at the highest current injection of 300 pA (Figure 2A, B, Supplemental Table 2). Consistently, the rheobase of WT overexpressing neurons was significantly increased (Figure 2C, Supplemental Table 2), their resting membrane potential was hyperpolarized (Figure 2D, Supplemental Table 2), and their input resistance and AP threshold were increased, whereas none of these parameters were changed in the R359C variant compared to control conditions (Supplemental Figure 2A and B, Supplemental Table 2). Peak amplitudes and AP half-width did not differ between the different conditions (Supplemental Figure 2C and D). Additionally, K_V_7.5-WT overexpression resulted in a significant increase in M-current amplitude, whereas the R359C variant caused a significant decrease, indicating that the transfected channel has indeed a dnLOF effect on the endogenous K_V_7 channels (Figure 2E and F, Supplemental Table 2).

**Figure 2.**
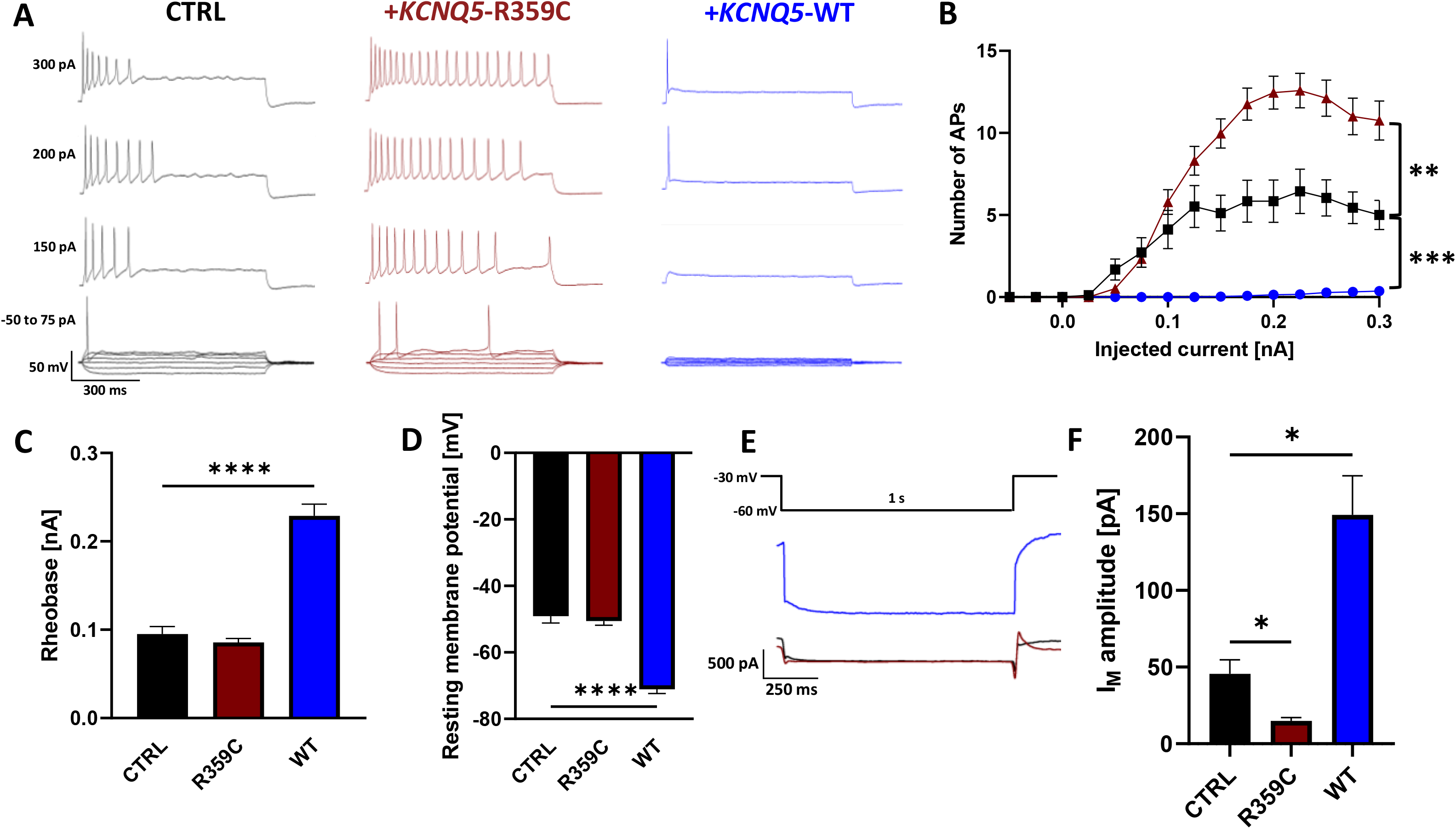
Effect of K_V_7.5-WT and -R359C expression in primary hippocampal neuronal culture. Neurons were either transfected with K_V_7.5-WT (blue) or –R359C (maroon), untransfected neurons were used as controls (CTRL; black). (**A**) Representative voltage traces of evoked action potentials at different current injections. (**B**) Number of evoked action potentials (APs) versus injected current. WT (blue dots) overexpression significantly reduces the area under the curve, whereas R359C (maroon triangles) transfection significantly increases it as compared to CTRL neurons (black squares). Shown are mean ± SEM. WT overexpression significantly increased rheobase (**C**) and decreased resting membrane potential (**D**). (**E**) Voltage protocol and representative current traces of the M-current. WT overexpression significantly increased M-current peak amplitude and significantly decreased it in R359C transfected neurons. * *p* < 0.05, ** *p* < 0.01; *** *p* < 0.001; **** *p* < 0.0001 (one-way ANOVA with Dunnett’s *post hoc* test or Kruskal-Wallis test Benjamini, Krieger and Yekutieli’s *post hoc* test). Statistical analysis and numbers of recorded cells are listed in Supplemental Figure 2.

### Pharmacological characterization of the K_V_7.5-R359C variant in transfected primary cultured hippocampal mouse neurons

Next, we investigated if the drugs previously tested in HEK cells also had effects on neuronal properties to restore neuronal dysfunction induced by the R359C variant. As WT overexpressing neurons had shown such a severe loss of firing activity in the previous set of experiments, drugs were not tested under these conditions, as an expected further decrease in firing would have been barely measurable. We therefore first investigated the effect of RTG on R359C-transfected and on untransfected neurons. RTG administration (10 µM) reduced neuronal firing in both conditions, however less strong as K_V_7.5-WT overexpression (Figure 3A). The input-output curves under RTG administration showed a flattening in both conditions and the areas under the curve were no longer significantly different between the R359C neurons treated with RTG and untransfected neurons without RTG, whereas RTG-treated untransfected neurons showed a significant decrease of the area under the curve (Figure 3B, Supplemental Table 2). RTG administration significantly increased rheobase and significantly decreased resting membrane potential in untransfected and R359C-transfected neurons (Figure 3C and D, Supplemental Table 2). Only untransfected neurons showed a significant decrease in input resistance and increase in AP threshold in the presence of RTG (Supplemental Figure 2E and F, Supplemental Table 2). AP peak amplitude and half-width were unaffected in both conditions with RTG compared to those without RTG (Supplemental Figure 2G and H). Furthermore, the M-current amplitude was significantly increased in R359C-transfected neurons by RTG, reaching comparable levels as untreated and untransfected control neurons (Figure 3E and F, Supplemental Table 2).

**Figure 3.**
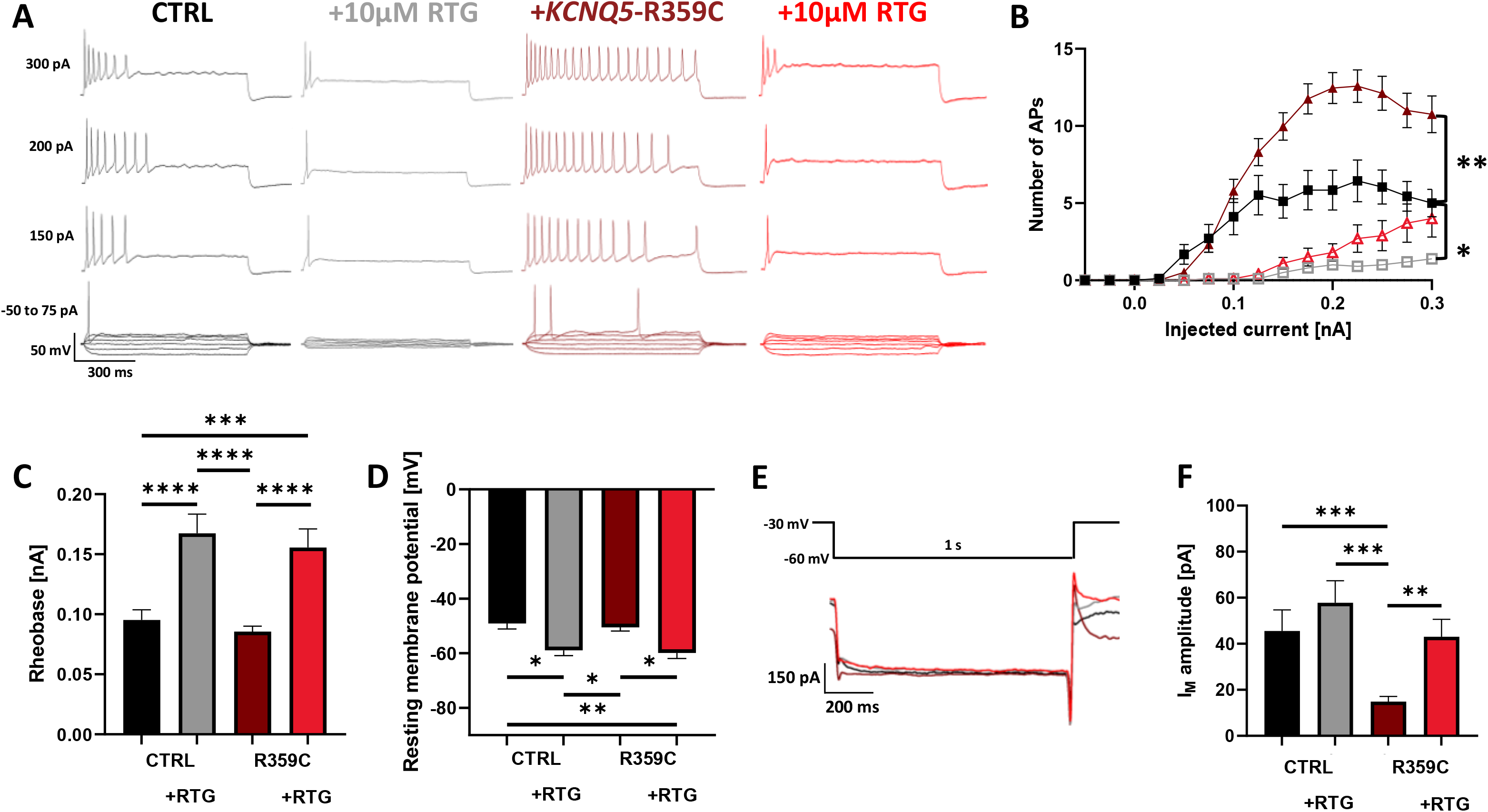
RTG counteracts K_V_7.5-R359C variant in transfected primary hippocampal neurons. (**A**) Representative voltage traces of evoked action potentials at different current injections in untreated CTRL neurons (black), RTG-treated CTRL neurons (grey), R359C expressing neurons (maroon) and RTG-treated R359C expressing neurons (red). (**B**) Number of evoked action potentials (APs) plotted against injected current. RTG treatment significantly decreases the area under the curve for R359C transfected (red outlined triangles) and CTRL neurons (grey outlined squares). Shown are mean ± SEM. Application of RTG significantly increased rheobase (**C**) and decreased resting membrane potential (**D**) in both conditions. (**E**) Voltage protocol and representative current traces of the M-current. RTG increased M-current peak amplitude in R359C transfected neurons back to untreated CTRL levels. * *p* < 0.05, ** *p* < 0.01; *** *p* < 0.001; **** *p* < 0.0001 (two-way ANOVA with Tukey’s *post hoc* test). Statistical analysis and numbers of recorded cells are listed in Supplemental Figure 2.

We next performed the same experiments under application of 100 µM GBP. While GBP did not show any significant effect in untransfected neurons on any of the investigated properties (Figure 4, Supplemental Table 2), in R359C-transfected neurons GBP significantly decreased the area under the curve of the input output curve, significantly increased the rheobase and M-current amplitude, reaching comparable levels as under control conditions (Figure 4B-F, Supplemental Table 2). None of the other parameters were significantly changed (Supplemental Figure 2I-L, Supplemental Table 2). Similar results were observed for the intracellular application of ZnCl_2_ which did not have a significant effect on untransfected neurons apart from significantly reducing the M-current (Figure 5, Supplemental Table 2). However, it showed an effect in the R359C-transfected neurons by decreasing the input-output area under the curve back to control levels, while significantly increasing the rheobase (Figure 5B and C, Supplemental Table 2). Additionally, a slight decrease in input resistance was observed compared to control conditions in untransfected neurons (Supplemental Figure 2M, Supplemental Table 2). Similarly, ZnCl_2_ application was able to increase the M-current amplitude enough to make it less significantly different as compared to untransfected cells, but was not significantly different when compared to R359C-transfected cells (Figure 5E and F, Supplemental Table 2). No other changes were observed (Supplemental Figure 2B-D, Supplemental Table 2). These results show that all three tested treatments can increase the neuronal M-current in neurons carrying a dnLOF variant and thereby decrease neuronal firing frequency, with RTG showing the strongest effect with the tested drug concentrations.

**Figure 4.**
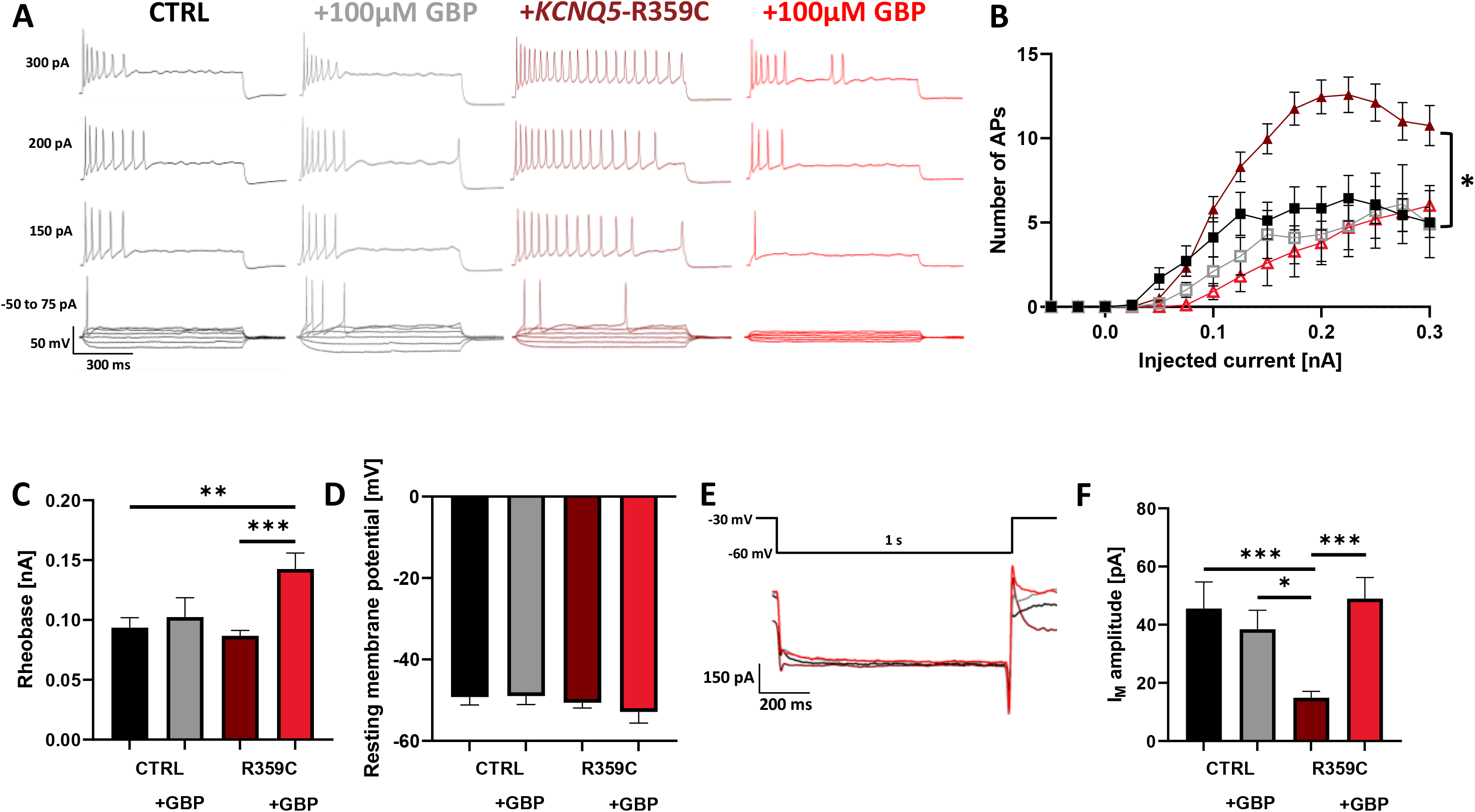
GBP has no effect on untransfected neurons but counteracts K_V_7.5-R359C variant in transfected primary hippocampal neurons. (**A**) Representative voltage traces of evoked action potentials at different current injections in untreated CTRL neurons (black), GBP-treated CTRL neurons (grey), R359C expressing neurons (maroon) and GBP-treated R359C expressing neurons (red). (**B**) Number of evoked action potentials (APs) versus injected current. GBP treatment significantly decreases the area under the curve for R359C transfected neurons (red outlined triangles). Shown are mean ± SEM. Application of GBP significantly increased rheobase (**C**) and has no effect on resting membrane potential (**D**) in R359C expressing neurons. (**E**) Voltage protocol and representative current traces of the M-current. GBP increased M-current peak amplitude in R359C transfected neurons back to untreated CTRL levels. * *p* < 0.05, ** *p* < 0.01; *** *p* < 0.001; **** *p* < 0.0001 (two-way ANOVA with Tukey’s *post hoc* test). Statistical analysis and numbers of recorded cells are listed in Supplemental Figure 2.

**Figure 5.**
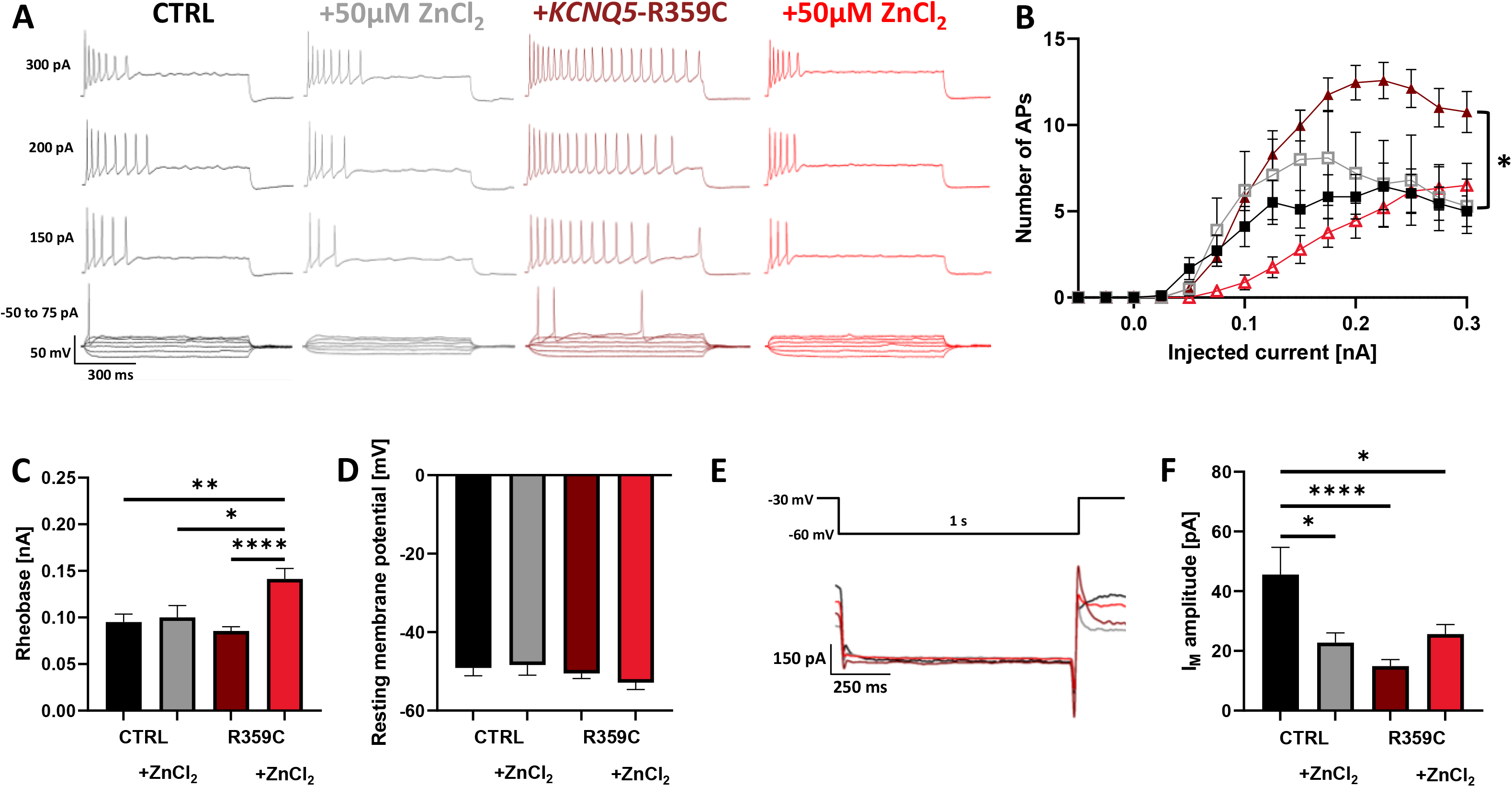
Intracellular application of ZnCl_2_ does not affect untransfected neurons but affects K_V_7.5-R359C variant in transfected primary hippocampal neurons. (**A**) Representative voltage traces of evoked action potentials at different current injections in untreated CTRL neurons (black), GBP-treated CTRL neurons (grey), R359C expressing neurons (maroon) and GBP-treated R359C expressing neurons (red). (**B**) Number of evoked action potentials (APs) plotted against injected current. Intracellular application of ZnCl_2_ significantly decreases the area under the curve for R359C transfected neurons (red outlined triangles). Shown are mean ± SEM. Intracellular application of ZnCl_2_ significantly increased rheobase (**C**) but has no effect on resting membrane potential (**D**) in R359C expressing neurons. (**E**) Voltage protocol and representative current traces of the M-current. While intracellular application of ZnCl_2_ decreased M-current peak amplitude in CTRL neurons, M-current peak amplitude was increased in R359C transfected neurons, yet they were still significantly decreased compared to untreated CTRL levels. * *p* < 0.05, ** *p* < 0.01; *** *p* < 0.001; **** *p* < 0.0001 (two-way ANOVA with Tukey’s *post hoc* test). Statistical analysis and numbers of recorded cells are listed in Supplemental Figure 2.

### Functional and pharmacological characterization of K_V_7.5-WT and -R359C variant expression on the medium afterhyperpolarization in transfected primary hippocampal mouse neurons

While recording the AP trains of neurons overexpressing K_V_7.5-WT channels, we observed a stark decrease of the membrane potential after an initial recovery from the afterhyperpolarization (Figure 2A). As the M-current is slowly activating and non-inactivating, this was perceived as a potential direct effect of the increased M-current in these neurons which is slowly activating after the AP and not inactivating until the end of the current step. To further investigate this effect, the resting membrane potential at the end of the train was normalized to the maximum resting membrane potential after the last AP. This was performed for the AP trains evoked at 200 pA, as WT overexpressing neurons would not fire APs prior to this step, and untransfected as well as R359C-transfected neurons would start to run into depolarization blocks at higher injected currents. This revealed that untransfected neurons underwent a slight decline in resting membrane potential after recovering from the last AHP of the AP train which was significantly increased in WT overexpressing neurons. On the contrary, R359C-transfected neurons did not show such a decline in resting membrane potential (Figure 6A, Supplemental Table 2). RTG application significantly increased this decline in untransfected neurons and initiated it in R359C neurons bringing them to the same level as untreated and untransfected neurons (Figure 6D, Supplemental Table 2). GBP did not have an effect in untransfected neurons but initiated a slight decline in R359C that was not significantly different from untreated and untransfected neurons (Figure 6G, Supplemental Table 2). Administration of ZnCl_2_ did not have an effect on either R359C transfected or untransfected neurons (Figure 6K, Supplemental Table 2). These results indicated a potential role of K_V_7.5 in the medium and slow AHP, which is consistent with previous studies identifying K_V_7.5 channels as contributors to the medium and slow AHP in the CA3 region of the mouse hippocampus (Tzingounis et al. 2010). Hence, the I_mAHP_ was recorded to examine the effect of a dnLOF or an overexpression of K_V_7.5 channels on this current. While neurons transfected with R359C showed a significant decrease in the amplitude of the medium AHP current, WT overexpression resulted in a non-significant increase (Figure 6B and C, Supplemental Table 2). RTG, GBP and intracellular application of ZnCl_2_ all increased R359C amplitudes so they were no longer significantly different from control levels, while untransfected neurons did not display significant changes by RTG or GBP treatment (Figure 6E, F, H, J, Supplemental Table 2). However, zinc application caused a significant reduction (Figure 6L and M, Supplemental Table 2). This indicates that the dnLOF variant reduces not only the M-current but in turn also decreases the I_mAHP_ explaining the significantly increased firing frequency observed in transfected neurons, while the opposite occurs in neurons overexpressing K_V_7.5 channels and with drug applications under conditions of reduced M-current by the dnLOF variant.

**Figure 6.**
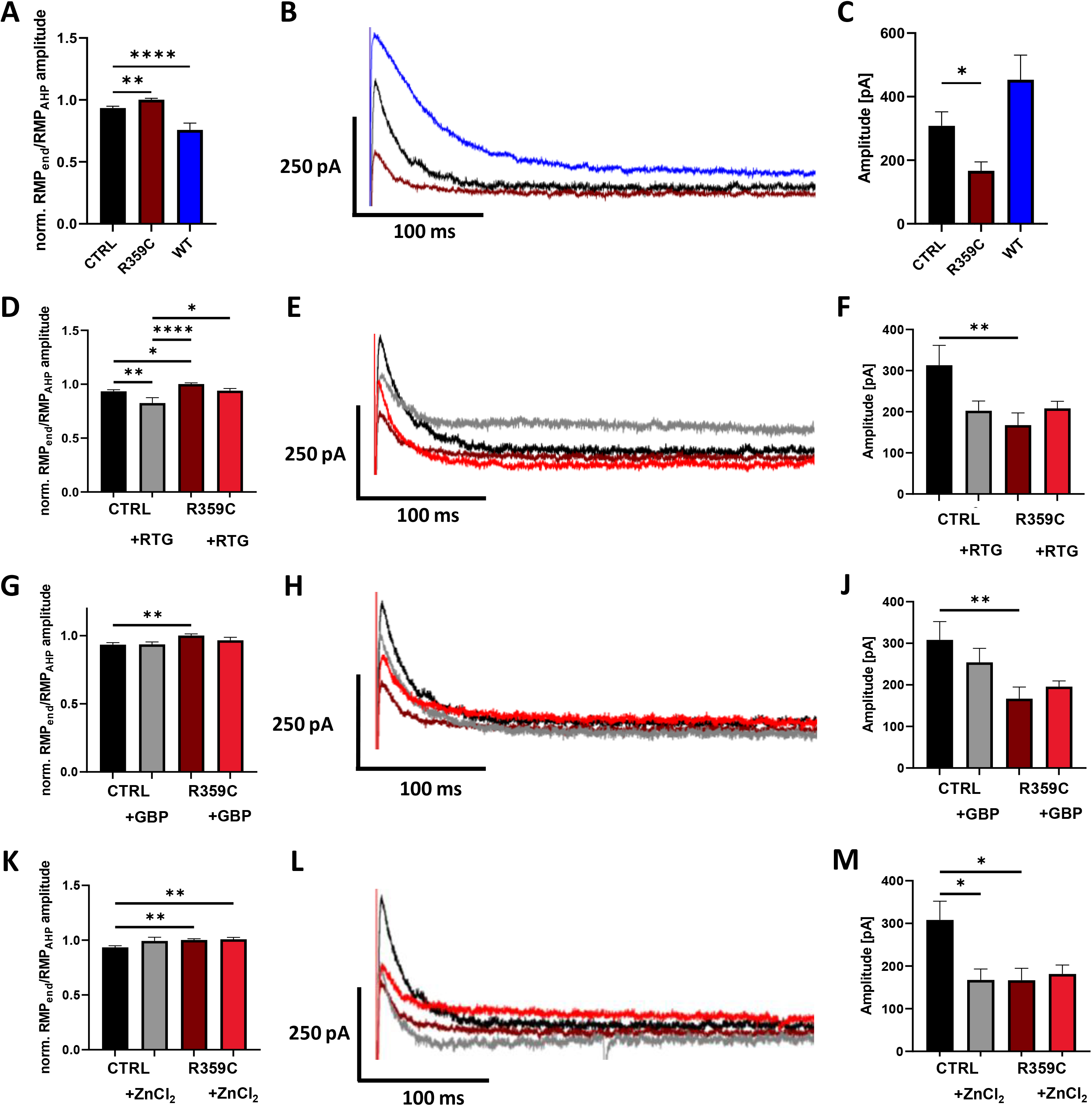
Expression of K_V_7.5-R359C decreases I_mAHP_ amplitude and is counteracted by RTG, GBP and intracellular application of ZnCl_2_ in primary hippocampal neuronal culture. (**A**) Overexpression of KV7.5-WT subunits significantly reduces resting membrane potential after an initial afterhyperpolarization of the last evoked AP, whereas the R359C does not show any reduction and is significantly increased as compared to the CTRL. RTG (**D**) significantly reduced normalized resting membrane potential amplitudes in CTRL and R359C transfected neurons, while GBP (**G**) only did so in the R359C expressing neurons. Intracellular application of ZnCl_2_ (**K**) did not have any effect. Middle panel displays representative I_mAHP_ traces for untreated conditions (**B**), under RTG (**E**), GBP (**H**) or intracellular ZnCl_2_ application (**L**). (**C**) K_V_7.5-R359C expression significantly reduces I_mAHP_ amplitude, while the increase in K_V_7.5-WT overexpressing neurons is not significant. RTG (**F**) and GBP (**J**) increase mAHP current amplitude in R359C transfected neurons. Intracellular application of ZnCl_2_ (**M**) increased R359C transfected I_mAHP_ amplitude just enough to make it no longer different from untransfected neurons, whereas it caused a significant decrease in untransfected neurons. * *p* < 0.05, ** *p* < 0.01; *** *p* < 0.001; **** *p* < 0.0001 (one-way ANOVA with Dunnett’s *post hoc* test, Kruskal-Wallis test Benjamini, Krieger and Yekutieli’s *post hoc* test, or two-way ANOVA with Tukey’s *post hoc* test). Statistical analysis and numbers of recorded cells are listed in Supplemental Figure 2.

## Discussion

Despite the discovery of many new effective antiepileptic drugs in the past decades, approximately 30% of epilepsy patients remain in a pharmacoresistant state (Picot et al. 2008, Fattorusso et al. 2021). Advances in next generation sequencing in the last three decades not only resulted in the discovery of many epilepsy-associated genes but also a better understanding of how the genotype and phenotype are correlated (Johannesen et al. 2022, reviewed in Oyrer et al. 2018). Knowing whether the phenotype is caused by a gain- or loss-of-function has proven to be crucial in opening doors to personalized precision medicine especially for pharmacoresistant patients (Hedrich et al. 2021, reviewed in Knowles et al. 2022). Here, we investigated the effect the dnLOF *KCNQ5*-/K_V_7.5-R359C variant – which we previously described in a GGE family including a pharmacoresistant patient (Krüger et al. 2022) – on neuronal firing and tested three different K_V_7 channel openers as potential treatment options *in vitro*.

Experiments in HEK cells showed that RTG, a well-established K_V_7 channel opener (Wuttke et al. 2005), was able to fully reverse the complete dnLOF of the R359C variant. GBP (100 µM) was also able to significantly increase current density and M-current in R359C homomeric and heteromeric channels although to a lesser degree than RTG making it a potential treatment option for K_V_7.5 LOF patients, as RTG was taken off the market due to severe side effects (Clark et al. 2015). These results further show that the variant is fully functional if enabled to open. Furthermore, intracellular application of zinc was tested as its proposed mechanism of action is to activate K_V_7 channels by decreasing their inhibition by calmodulin and reducing their dependence on PIP_2_ for opening (Yang et al. 2023, Gao et al. 2017), potentially counteracting the previously found altered PIP_2_ binding of the R359C variant. However, intracellular application of zinc only showed slight effects in R359C homomeric channels whereas it reached results comparable to GBP in heteromeric channels. This indicates that the effect of the variant cannot be overcome by zinc and the effect seen in the heteromeric channels might be due to its effect on WT subunits.

Expression of the R359C variant in hippocampal neuronal cultures significantly increased neuronal firing by reducing the M-current and thereby reducing the medium afterhyperpolarization. Overexpression of K_V_7.5-WT channels caused the opposite effect and practically silenced neurons by increasing the medium afterhyperpolarization through amplifying the M-current. Usually this mechanism is crucial for intrinsic control of excitability, for example by enabling burst firing through acetylcholine release (Brown and Adams 1980). Our findings underline the delicate balance neurons need to maintain, as both a reduction and an amplification in M-current can have detrimental effects on a single neuron and the adjacent network. Interestingly, LOF as well as GOF variants in K_V_7.5 have been linked to epileptic phenotypes and/or intellectual disability. While a LOF seems to induce milder phenotypes, such as GGE, with or without intellectual disability, GOF variants have been linked to severe phenotypes such as DEE (Lehman et al. 2017, Rosti et al. 2019, Krüger et al. 2022, Wei et al. 2022, Nappi et al. 2022). It remains unclear how exactly alterations in both directions can generate an epileptic phenotype and further studies should be conducted in animal models to uncover the underlying mechanisms on a network level. Here, we were able to counteract the effect of the variant by RTG or GBP application, which both reduced firing frequency by increasing the M-current and medium AHP current. Hence, GBP as a licensed anti-seizure medication could be tried in pharmacoresistant patients carrying LOF *KCNQ5* variants. Interestingly, intracellular application of zinc did not significantly alter untransfected neurons apart from decreasing their M-current. In contrast, zinc reduced firing frequency in R359C-transfected neurons back to the level of untransfected neurons. As zinc can also modulate many other currents, such as the A- and K-current at the administered concentration (Easaw et al. 1999), the observed effect is likely due to altering these channels’ properties as well.

In summary, our data provide evidence that the R359C causes the epileptic phenotype in the patients by decreasing the M-current and slow AHP current significantly, which results in amplified neuronal firing while WT overexpression has the reverse effect and nearly abolishes firing. RTG and GBP were both able to recover neuronal firing rates back to levels in untransfected neurons by increasing the M-current as well as slow AHP current. As RTG is currently not available as a treatment option to patients, we propose GBP as a treatment option for patients carrying *KCNQ5* LOF variants.

## Authors contributions

JK and HL designed the study and experiments. JK performed experiments and analysed data. JK and HL interpreted data. JK and HL wrote the manuscript. Both authors read, revised, and approved the manuscript.

## Supplemental Figures

**Supplemental Figure 1. Pharmacological characterization of heteromeric K_V_7.5-WT and -R359C channels in HEK cells.** HEK cells were transfected with either KCNQ5-WT or KCNQ5-WT- and -R359C DNA and recorded under either RTG (left panel), GBP (middle panel) or ZnCl_2_ application (right panel). (**A**) Representative K^+^ current traces at +60 mV for K_V_7.5-WT (control black, drug treated grey) and K_V_7.5-WT and -R359C (control maroon, drug treated red). (**B-D**) Peak K+ currents were normalized to cell capacitances and plotted versus voltage. WT+R359C (filled marron triangles) currents are significantly decreased as compared to WT (filled black squares) currents showing no difference to CTRL cells (green crosses). RTG increased WT currents, yet not significantly (grey outlined squares), and increased R359C peak current density (red outlined triangles) back to WT levels. GBP had no significant effect on WT currents but significantly increased mutant currents, though they were still significantly decreased as compared to WT. Intracellular application of ZnCl_2_ did not significantly increase currents in the WT or mutant channels. (**E-G**) Maximum current peak density comparison at +60 mV of WT (black) and WT+R359C (maroon) to drug treatment (grey or red, respectively). (**H-J**) Voltage dependent activation curves. Normalized tail currents were fit with a Boltzmann function. No such relationship could be established for the WT+R359C as currents were too small. RTG significantly left-shifted R359C as compared to WT while WT slope was significantly increased by zinc. (**K-M**) Representative M-current traces and voltage protocol. (**N-P**) M-current peak was significantly increased in WT+R359C by RTG, GBP, and ZnCl_2_ application. Shown are means ± SEM (**E-G** and **N-P**). * *p* < 0.05, ** *p* < 0.01; *** *p* < 0.001; **** *p* < 0.0001 (one-way ANOVA with Dunnett’s *post hoc* test or Kruskal-Wallis test with Benjamini, Krieger and Yekutieli’s *post hoc* test). Supplemental Table 1 provides exact numbers and statistical analysis.

**Supplemental Figure 2. Effect of K_V_7.5-WT and -R359C expression on input resistance, threshold, peak amplitude and action potential half-width in untreated and drug-treated primary hippocampal neuronal culture.** K_V_7.5-WT expression neurons present significantly decreased input resistance (A) and significantly increased threshold (B) as compared to CTRL. RTG application had a similar effect on CTRL but not R359C neurons (E, F). GBP application did not have a significant effect on input resistance (I) and threshold (J). Intracellular application of ZnCl_2_ significantly decreased input resistance in R359C expressing neurons (M), but had no effect on action potential threshold (N). Action potential peak amplitude (C, G, K, O) and half-width (D, H, L, P) were not affected in any condition. * *p* < 0.05, ** *p* < 0.01; *** *p* < 0.001; **** *p* < 0.0001 (one-way ANOVA with Dunnett’s *post hoc* test or Kruskal-Wallis test Benjamini, Krieger and Yekutieli’s *post hoc* test). Statistical analysis and numbers of recorded cells are listed in Supplemental Figure 2.

## Supplemental Tables

**Supplemental Table 1.** Summary of biophysical properties of K_v_7.5-WT and -R359C channels under control conditions, RTG, GBP, or intracellular zinc application in homomeric and heteromeric state. Data are displayed as mean ± SEM; * *p* < 0.05, ** *p* < 0.01; *** *p* < 0.001; **** *p* < 0.0001 as compared to WT (one-way ANOVA with Dunnett’s *post hoc* test,or Kruskal-Wallis test with Benjamini, Krieger and Yekutieli’s *post hoc* test).

|  | Homomeric expression |  |  |  |  |  |  | Heteromeric expression |  |  |  |  |  |  |
| --- | --- | --- | --- | --- | --- | --- | --- | --- | --- | --- | --- | --- | --- | --- |
|  |  |  | Activation kinetics |  |  |  |  |  |  | Activation kinetics |  |  |  |  |
| | Peak current<br>density<br>[pA/pF] | n | $V_{1/2}$<br>[mV] | k-slope | n | $I_M$ [pA] | n | Peak current<br>density [pA/pF] | n | $V_{1/2}$<br>[mV] | k-slope | n | $I_M$ [pA] | n |
| WT | $273.59 \pm 51.77$ | 16 | -28.26<br>$\pm 4.13$ | $16.66 \pm 2.35$ | 14 | $84.78 \pm 17.64$ | 16 | $327.72 \pm 72.24$ | 11 | -31.45<br>$\pm 3.46$ | $12.20 \pm 2.40$ | 10 | $70.82 \pm 15.47$ | 9 |
| + RTG | $814.76 \pm 132.61$ | 10 | -33.64<br>$\pm 3.00$ | $16.48 \pm 1.87$ | 10 | $297.64 \pm 73.78$ | 10 | $700.87 \pm 98.61$ | 10 | -33.86<br>$\pm 3.74$ | $17.98 \pm 2.65$ | 10 | $256.89 \pm 53.28$ | 9 |
| + GBP | $169.95 \pm 20.87$ | 10 | -37.57<br>$\pm 2.28$ | $10.60 \pm 2.30$ | 10 | $74.58 \pm 13.52$ | 10 | $176.52 \pm 19.72$ | 7 | -37.14<br>$\pm 7.20$ | $14.50 \pm 4.23$ | 7 | $52.43 \pm 13.59$ | 10 |
| + zinc | $525.56 \pm 246.01$ | 10 | -26.97<br>$\pm 4.12$ | $15.82 \pm 2.91$ | 10 | $159.34 \pm 49.37$ | 10 | $501.11 \pm 125.54$ | 10 | -20.66<br>$\pm 6.62$ | $21.26 \pm 3.62^*$ | 10 | $179.36 \pm 23.59$ | 8 |
| R359C | $18.47 \pm 5.47$ | 10 | - | - | - | $5.86 \pm 1.35$ | 10 | $38.28 \pm 4.95$ | 10 | - | - | - | $6.56 \pm 2.11$ | 10 |
| + RTG | $352.49 \pm 56.59$ | 10 | -47.49<br>$\pm 5.84^{**}$ | $28.95 \pm 3.78^{**}$ | 10 | $66.81 \pm 6.61$ | 10 | $413.98 \pm 72.10$ | 10 | -45.85<br>$\pm 3.61^{**}$ | $21.07 \pm 3.88$ | 10 | $130.55 \pm 27.03$ | 10 |
| + GBP | $91.56 \pm 14.21$ | 9 | -37.21<br>$\pm 2.57$ | $16.62 \pm 2.81$ | 9 | $18.32 \pm 2.16$ | 10 | $81.24 \pm 13.90$ | 10 | -37.57<br>$\pm 4.81$ | $16.69 \pm 1.70$ | 9 | $20.20 \pm 1.89$ | 10 |
| + zinc | 39.58 ± 8.56 | 10 | -27.25<br>± 9.05 | 17.11 ±<br>4.80 | 6 | 10.1 ± 5.76 | 9 | 104.64 ± 58.13 | 10 | -37.89<br>± 3.09 | 16.67 ±<br>2.69 | 8 | 45.20 ± 22.87 | 10 |
| CTRL | 28.86 ± 1.76 | 10 | - | - | - | 2.63 ± 1.36 | 10 | 28.86 ± 1.76 | 10 | - | - | - | 2.63 ± 1.36 | 10 |

**Supplemental Table 2.**
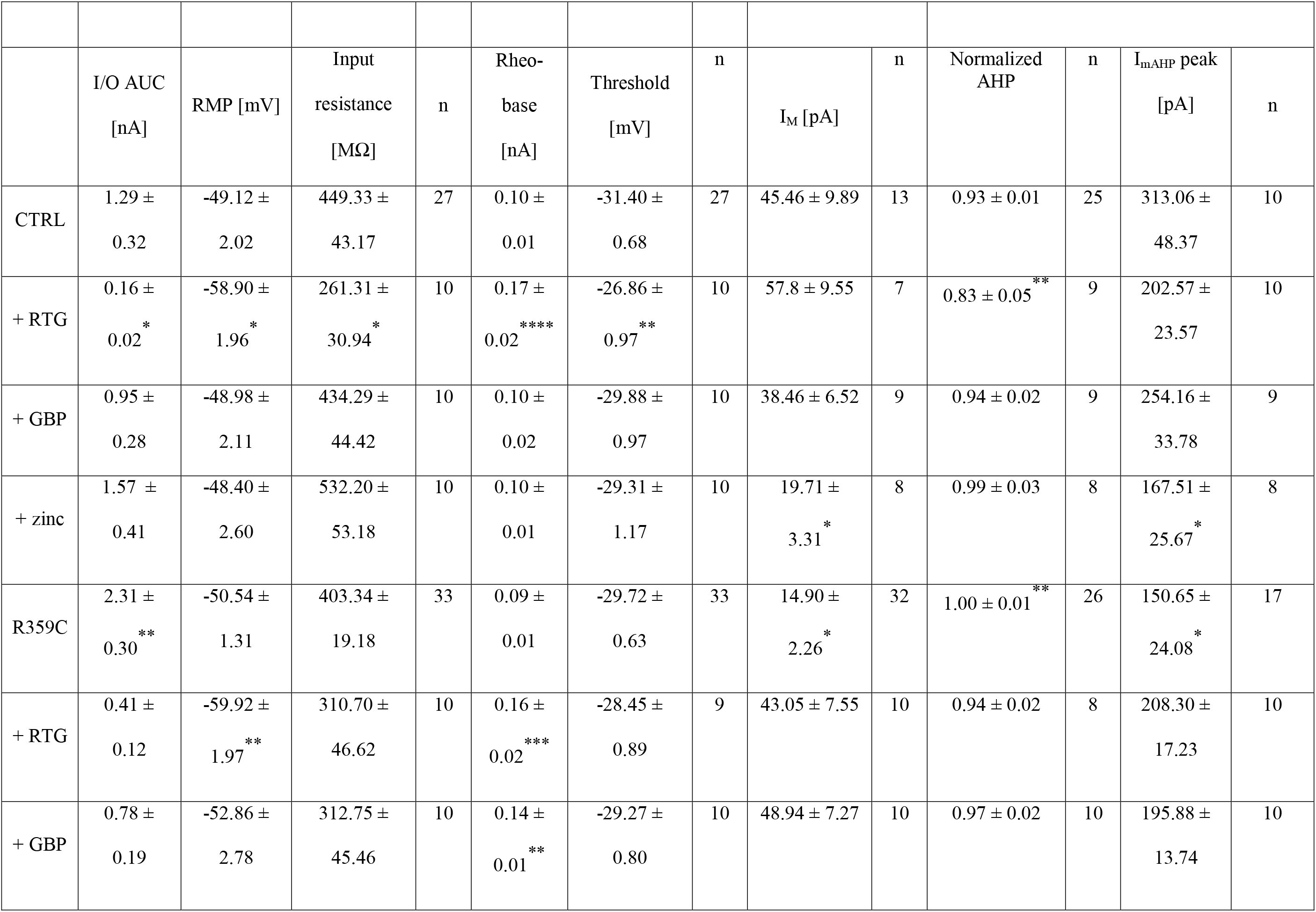
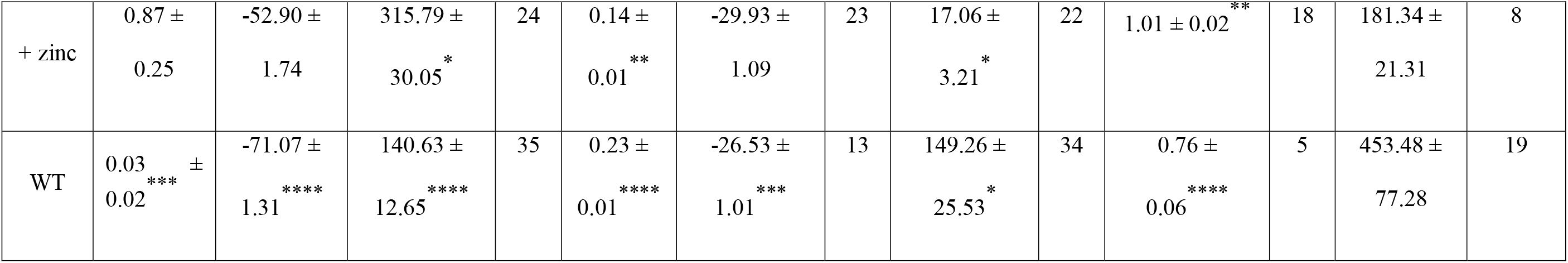
Summary of intrinsic neuronal properties in control, and K_v_7.5-WT or -R359C transfected hippocampal neurons under control conditions, RTG, GBP, or intracellular zinc application. Single action potential parameters were not significantly changed in any condition and are hence not listed. I/O AUC = input-output area under the curve; RMP = resting membrane potential. Data are displayed as mean ± SEM; * *p* < 0.05, ** *p* < 0.01; *** *p* < 0.001; **** *p* < 0.0001 (one-way ANOVA with Dunnett’s *post hoc* test, Kruskal-Wallis test with Benjamini, Krieger and Yekutieli’s *post hoc* test, or two-way ANOVA with Tukey’s *post hoc* test).

|  | I/O AUC<br>[nA] | RMP [mV] | Input<br>resistance<br>[MΩ] | n | Rheo-<br>base<br>[nA] | Threshold<br>[mV] | n | I<sub>M</sub> [pA] | n | Normalized<br>AHP | n | I<sub>mAHP</sub> peak<br>[pA] | n |
| --- | --- | --- | --- | --- | --- | --- | --- | --- | --- | --- | --- | --- | --- |
| CTRL | 1.29 ±<br>0.32 | -49.12 ±<br>2.02 | 449.33 ±<br>43.17 | 27 | 0.10 ±<br>0.01 | -31.40 ±<br>0.68 | 27 | 45.46 ± 9.89 | 13 | 0.93 ± 0.01 | 25 | 313.06 ±<br>48.37 | 10 |
| + RTG | 0.16 ±<br>0.02 * | -58.90 ±<br>1.96 * | 261.31 ±<br>30.94 * | 10 | 0.17 ±<br>0.02 **** | -26.86 ±<br>0.97 ** | 10 | 57.8 ± 9.55 | 7 | 0.83 ± 0.05 ** | 9 | 202.57 ±<br>23.57 | 10 |
| + GBP | 0.95 ±<br>0.28 | -48.98 ±<br>2.11 | 434.29 ±<br>44.42 | 10 | 0.10 ±<br>0.02 | -29.88 ±<br>0.97 | 10 | 38.46 ± 6.52 | 9 | 0.94 ± 0.02 | 9 | 254.16 ±<br>33.78 | 9 |
| + zinc | 1.57 ±<br>0.41 | -48.40 ±<br>2.60 | 532.20 ±<br>53.18 | 10 | 0.10 ±<br>0.01 | -29.31 ±<br>1.17 | 10 | 19.71 ±<br>3.31 * | 8 | 0.99 ± 0.03 | 8 | 167.51 ±<br>25.67 * | 8 |
| R359C | 2.31 ±<br>0.30 ** | -50.54 ±<br>1.31 | 403.34 ±<br>19.18 | 33 | 0.09 ±<br>0.01 | -29.72 ±<br>0.63 | 33 | 14.90 ±<br>2.26 * | 32 | 1.00 ± 0.01 ** | 26 | 150.65 ±<br>24.08 * | 17 |
| + RTG | 0.41 ±<br>0.12 | -59.92 ±<br>1.97 ** | 310.70 ±<br>46.62 | 10 | 0.16 ±<br>0.02 *** | -28.45 ±<br>0.89 | 9 | 43.05 ± 7.55 | 10 | 0.94 ± 0.02 | 8 | 208.30 ±<br>17.23 | 10 |
| + GBP | 0.78 ±<br>0.19 | -52.86 ±<br>2.78 | 312.75 ±<br>45.46 | 10 | 0.14 ±<br>0.01 ** | -29.27 ±<br>0.80 | 10 | 48.94 ± 7.27 | 10 | 0.97 ± 0.02 | 10 | 195.88 ±<br>13.74 | 10 |
| + zinc | 0.87 ±<br>0.25 | -52.90 ±<br>1.74 | 315.79 ±<br>30.05 * | 24 | 0.14 ±<br>0.01 ** | -29.93 ±<br>1.09 | 23 | 17.06 ±<br>3.21 * | 22 | 1.01 ± 0.02 ** | 18 | 181.34 ±<br>21.31 | 8 |
| WT | 0.03 ±<br>0.02 *** | -71.07 ±<br>1.31 **** | 140.63 ±<br>12.65 **** | 35 | 0.23 ±<br>0.01 **** | -26.53 ±<br>1.01 *** | 13 | 149.26 ±<br>25.53 * | 34 | 0.76 ±<br>0.06 **** | 5 | 453.48 ±<br>77.28 | 19 |

