## Supplementary figures and images for "Retigabine and gabapentin restore channel function and neuronal firing of an epilepsy-associated dominant-negative *KCNQ5* variant"

### Supplemental Figure 1

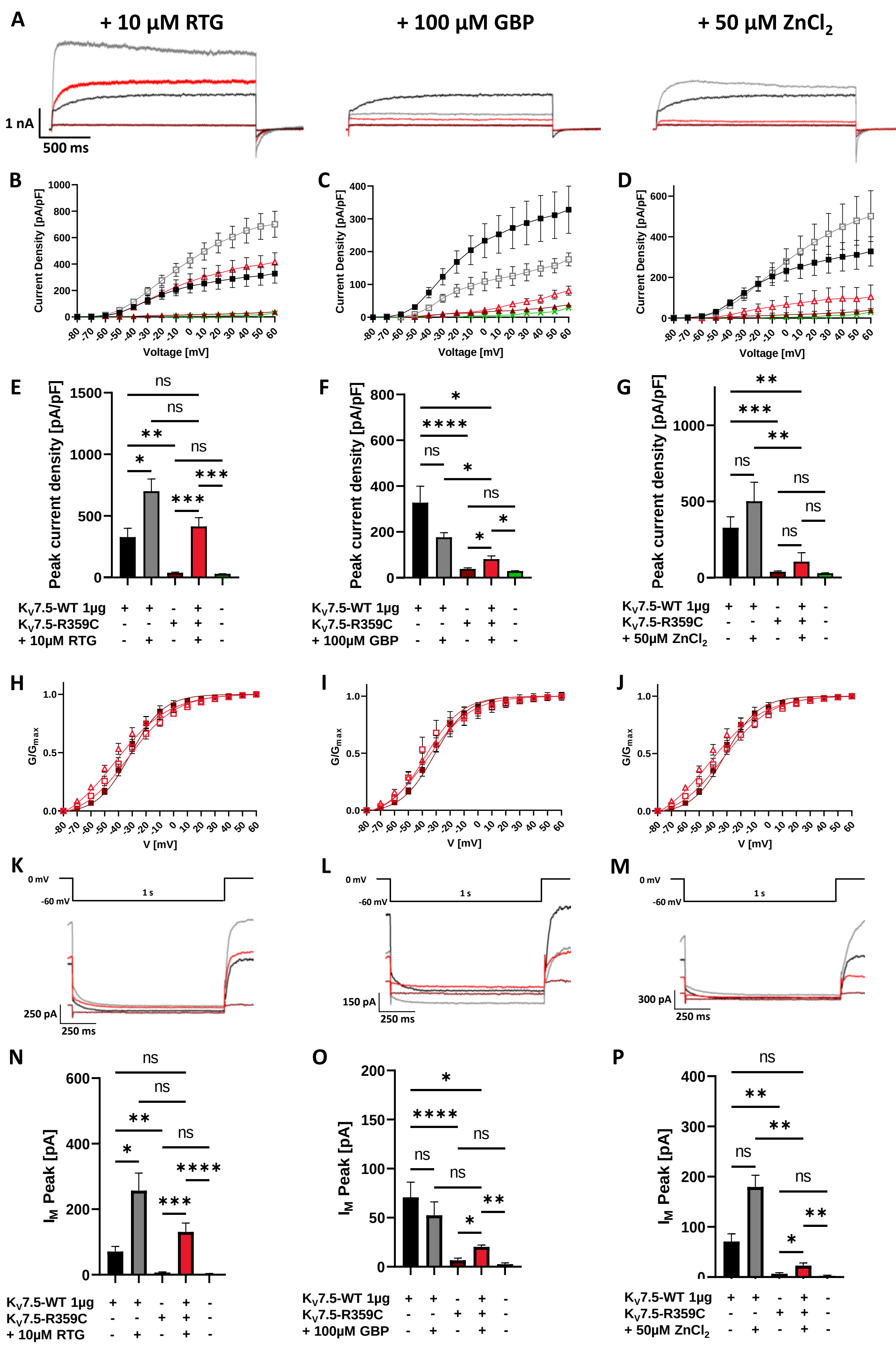

### Supplemental Figure 2

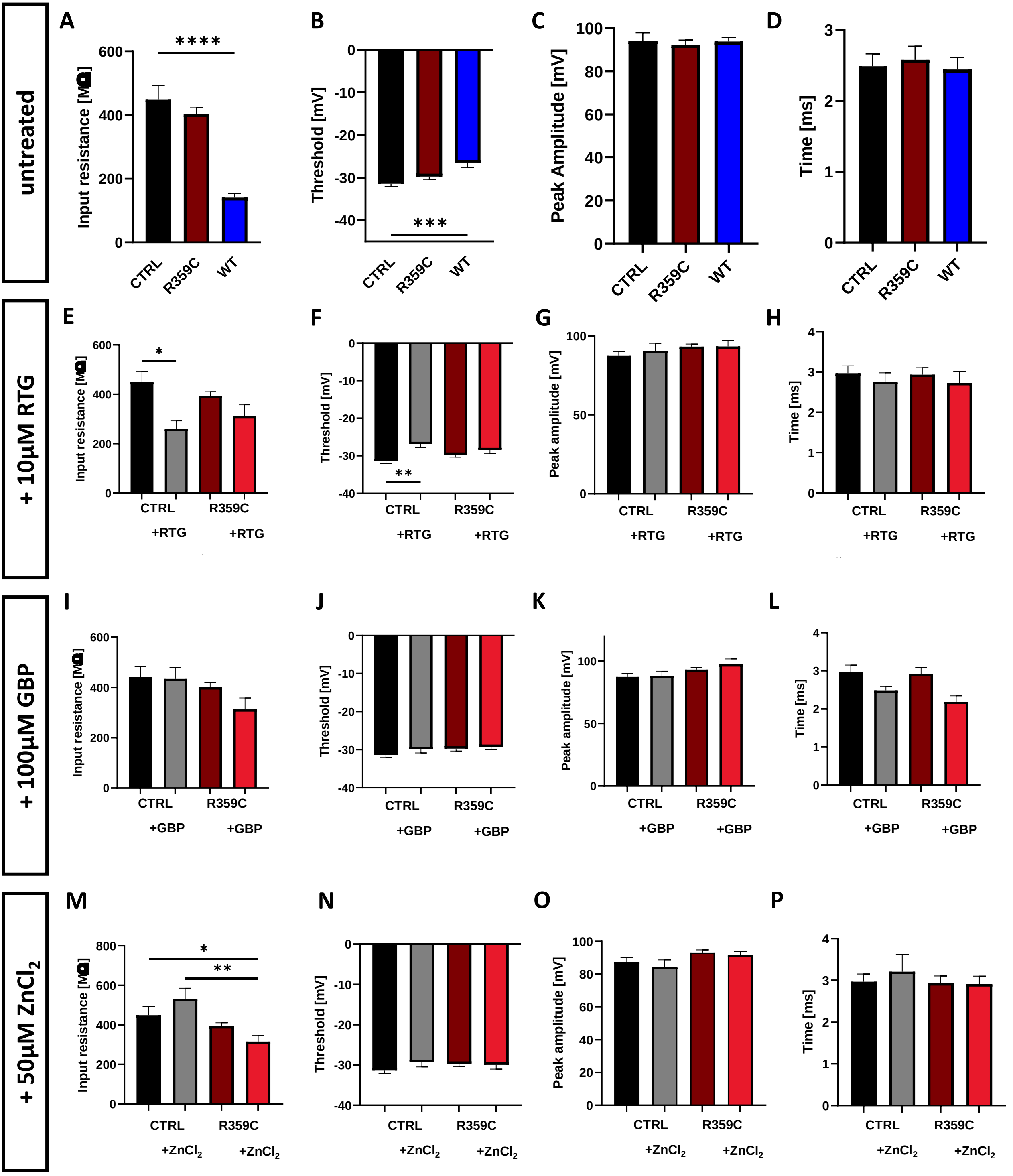
